# Proximity of astrocyte leaflets to the synapse determines memory strength

**DOI:** 10.1101/2022.01.30.478393

**Authors:** Aina Badia-Soteras, Tim S Heistek, Mandy S J Kater, Adrian Negrean, Huibert D Mansvelder, Baljit S Khakh, Rogier Min, August B Smit, Mark H G Verheijen

## Abstract

**Summary:** Astrocyte distal processes, known as leaflets or perisynaptic astrocyte processes (PAPs), fine-tune synaptic activity by clearing neurotransmitters and limiting extrasynaptic glutamate diffusion. While learning and memory depends on orchestrated synaptic activity of neuronal ensembles within the hippocampus, it is becoming increasingly evident that astrocytes residing in the environment of these synapses play a central role in shaping memories. However, how astroglial synaptic coverage contributes to mnemonic processing remains largely unknown. Here, we targeted astrocyte leaflet structure *in vivo* by depleting Ezrin, an integral leaflet-structural protein, in astrocytes of the adult hippocampal CA1 using a CRISPR-Cas9 genetic approach. This resulted in significantly smaller astrocyte territories and reduced astroglial synaptic coverage. In addition, using genetically encoded glutamate sensors and whole-cell patch-clamp recordings from pyramidal neurons, we found that Ezrin deletion and the resultant manipulation of leaflet structure boosted extrasynaptic glutamate diffusion and NMDA-receptor activation. Importantly, these cellular phenotypes translated to enhanced fear memory expression that was accompanied by increased activation of CA1 pyramidal neurons in the days after learning occurred. We show that Ezrin is critical for astrocyte morphology as well as for adult hippocampal synapse integrity and function. Our data show that astrocyte leaflet structure gates memory strength by regulating glutamate spillover in the vicinity of memory-related synaptic activity.

## Introduction

Astrocytes are morphologically complex cells that tile the brain with their extensive, ramified and specialized processes. A single astrocyte in the hippocampal *stratum radiatum* of mice can envelope ∽140,000 synapses in its territory (Bushong et al., 2002). These glial cells contact synapses with their thinnest terminal processes, referred to as leaflets or perisynaptic astrocyte processes (PAPs), which together with pre– and postsynaptic neuronal elements (Araque et al., 2014; Khakh and Sofroniew, 2015; Semyanov and Verkhratsky, 2021) are an integral feature of synapses throughout the CNS. Astrocytic leaflets fine-tune local synaptic transmission and plasticity because they are ideally positioned proximally to the synaptic cleft (Ghézali et al., 2016) and express molecular machinery that can modulate the extracellular space around synapses (Araque et al., 1999; Bazargani and Attwell, 2016).

Interestingly, neuronal activity has been reported to induce remodelling of astrocytic leaflets in the amygdala (Ostroff et al., 2014), somatosensory cortex (Henneberger et al., 2020) and the hypothalamus (Oliet, 2001; Panatier and Oliet, 2006) *in vivo*, and also in acute hippocampal slices (Bernardinelli et al., 2014; Henneberger et al., 2020; Lushnikova et al., 2009; Perez-Alvarez et al., 2014; Wenzel et al., 1991). Consistent with these findings, astrocyte leaflets are enriched in actin-associated proteins of the Ezrin Radixin Moesin (ERM) family that dynamically regulate membrane cytoskeleton linkage (Derouiche and Frotscher, 2001). In accord, Ezrin was found to be essential for astrocytic filopodia formation in culture (Lavialle et al., 2011) and for astrocyte morphogenesis in the developing somatosensory cortex (Zhou et al., 2019). However, whether Ezrin is required for leaflet structure and consequently for the integrity and function of synapses in the adult hippocampus, has remained unexplored.

Recent studies have used diverse approaches, such as chemogenetic activation of CA1 astrocytes (Adamsky et al., 2018), interference with astrocyte-neuron lactate transport (Gao et al., 2016; Suzuki et al., 2011), and genetic deletion of the gap junction protein Connexin 30 (Pannasch et al., 2014), to establish that astrocyte-neuron communication is involved in hippocampal-dependent memories. Despite these recent insights, the precise functional role of astrocytic leaflet structure in memory processing remains unclear. To gain insight into the role of astrocytic leaflets in synaptic function and memory, we developed a CRISPR-Cas9 viral approach to specifically delete Ezrin and thereby target astrocytic leaflet structure in the adult hippocampus. We found that the deletion of Ezrin in CA1 astrocytes resulted in smaller astrocyte territories and reduced astroglial coverage of excitatory synapses. This structurally compromised astrocyte-synapse interaction led to a decrease in synaptic glutamate and increased extrasynaptic glutamate diffusion as measured by whole-cell patch-clamp recordings of pyramidal neurons, and two-photon imaging of genetically encoded glutamate sensors, respectively. Moreover, these synaptic alterations had a significant consequential impact on contextual fear memory, resulting in enhanced retrieval that was accompanied by increased activation of the underlying supporting neuronal network. The present study thus reveals that the apposition of astrocyte leaflets to the synaptic cleft influences contextual fear memory expression and gates neuronal activation in a context-dependent manner.

## Results

### Structural manipulation of astrocyte leaflets by CRISPR-Cas9-mediated depletion of Ezrin

To interfere with leaflet structure *in vivo*, we specifically targeted Ezrin in CA1 astrocytes of adult mice by delivering an adeno-associated viral vector (AAV2/5) encoding the saCas9 enzyme and a single guide RNA (sgRNA) complementary to the Ezrin gene (exon-1), under the control of the minimal astrocyte-specific glial fibrillary acidic protein (GFAP) promoter (AAV2/5-GfaABC_1_D::Cas9-HA-Ezr) (Figure 1a). Stereotaxic delivery of this vector was restricted to dorsal CA1 astrocytes, with high penetrance (82.7 ± 2.9%; mean ± SEM; of astrocytes expressed Cas9) and very high specificity (Figure 1b,c). Next, we examined Ezrin levels by combining RNA scope *in situ* hybridization and immunohistochemistry (IHC) analysis, 4-5 weeks after AAV delivery. We found strong reduction (72%) of Ezrin RNA levels in the dorsal CA1 of mice expressing saCas9 together with the sgRNA against Ezrin (Ezr-saCas9) compared to controls expressing only Cas9 (Control: 27.4 ± 2.7, Ezr-saCas9: 7.5 ± 1.2 RNA molecules per astrocyte) (Figure 1d,e). Similarly, Ezrin protein levels were found to be reduced in Ezr-saCas9 mice compared to controls (100 ± 11.6%, Ezr-saCas9:27.1 ± 5.7% Ezrin signal a.u relative to control) (Figure 1f,g). To determine possible virus-induced reactive astrogliosis (Ortinski et al., 2010) we investigated the number of astrocytes (Control: 100 ± 9.5%, Ezr-saCas9: 110 ± 4.4% GFAP^+^ astrocytes relative to control), and GFAP protein levels (Control: 100 ± 1.2%, Ezr-saCas9: 88.7 ± 4.8% GFAP signal a.u relative to control), but did not find major signs of gliosis (Figure S1a-c). Thus, saCas9 enabled us to specifically target astrocytes in the CA1 region of the adult hippocampus, resulting in Ezrin protein level reduction by 73% within a five-week period.

**Figure 1.**
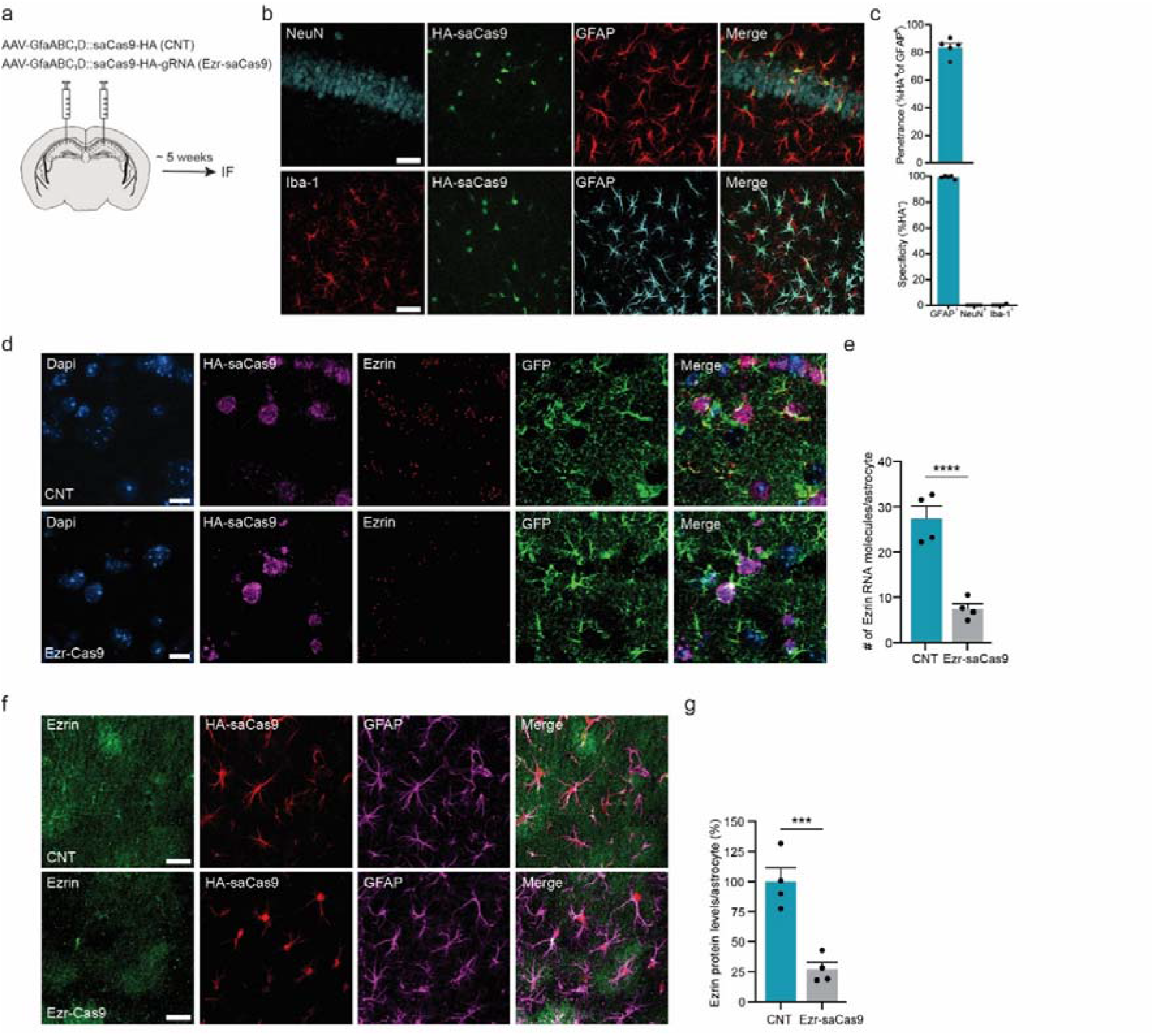
CRISPR-saCas9 viral approach to target Ezrin in adult hippocampal astrocytes. **a)** Schematic of experimental workflow. **b)** Representative images of the penetrance and specificity of viral vectors assessed by immuno-fluorescence (IF). Scale bar: 50 μm. **c)** Summary data of penetrance (718 cells from 5 mice) and specificity (NeuN: 837 cells from 5 mice, Iba-1: 364 cells from 5 mice). **d)** RNA scope *in situ* hybridization representative images for control and Ezr-saCas9 mice. Scale bar: 10 μm. **e)** Summary data of the number of RNA molecules of Ezrin per astrocyte (Control: 128 cells from 4 mice, Ezr-saCas9: 179 cells from 4 mice). Nested t test (t_305_ = 23.69, **** p <0.00001). **f)** IF representative images for control and Ezr-saCas9 mice. Scale bar: 30 μm. **g)** Summary data of Ezrin protein levels per astrocyte (%) (Control: 110 astrocytes from 4 mice, Ezr-saCas9: 102 astrocytes from 4 mice). Nested t test (t_82_ = 15.09, *** p = 0.0001). Data are presented as mean ± SEM.

### Astrocyte complexity and astrocyte-neuron proximity are decreased in the hippocampus of Ezr-saCas9 mice

Next, we determined how Ezrin shapes astrocyte morphology and leaflet structure in the adult hippocampus. To this end, we made use of a Napa-a viral vector (Octeau et al., 2018) to visualize astrocyte territories in the CA1 region of the hippocampus. We found that Ezr-saCas9 mice had smaller astrocyte territories compared to controls (Control: 1873 ± 69, Ezr-saCas9: 1631 ± 484 μm^2^) (Figure 2a-c). To determine whether this smaller territory in Ezr-saCas9 mice was accompanied by shorter astrocyte leaflets, we measured their proximity to synapses using the FRET-based Napa technique that captures spatial interactions within ∽10 nm range between astrocyte processes and synapses in acute brain slices (Octeau et al., 2018). Astrocytic membranes in the CA1 *stratum radiatum* were labelled with the FRET donor GFP (Napa-a) and Schaffer Collateral presynaptic terminals were labelled with the FRET acceptor mCherry (Napa-n), to allow detection of FRET signals at the synaptic scale. We prepared acute hippocampal slices 4-5 weeks after viral injection, and using confocal imaging to detect FRET signals in the CA1 (Figure 2d,e) we found that the number of synapses contacted by an astrocyte was significantly reduced in Ezr-saCas9 mice compared to control (Control: 26.3 ± 2.0, Ezr-saCas9: 19.9 ± 1.4 FRET ROIs/astrocyte) (Figure 2f,g). Importantly, this reduction in FRET was not due to differential expression of Napa-a or Napa-n across groups (Control: 1142 ± 101, Ezr-saCas9: 968.7 ± 160.9 Napa-a signal a.u; Control: 279.5 ± 54.1, Ezr-saCas9: 332.2 ± 53.2 Napa-n signal a.u) (Figure S2).

**Figure 2.**
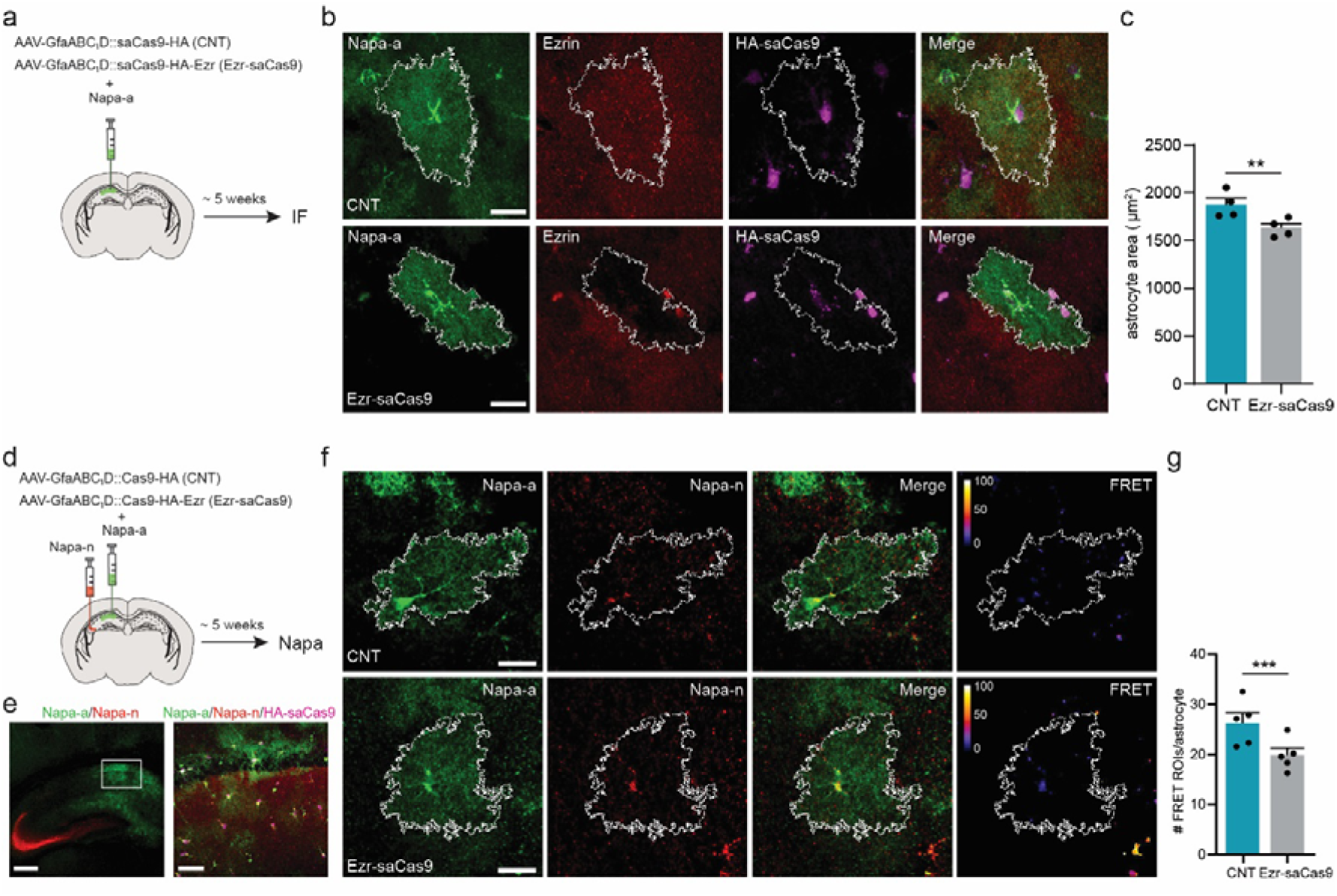
Deletion of Ezrin in mature astrocytes reduces morphological complexity and neuron-astrocyte interaction. **a)** Schematic of the experimental workflow to assess astrocyte morphology. **b)** Representative images of the astrocyte domain for control and Ezr-saCas9 mice. Astrocyte territory is outlined in white. Scale bar: 20 μm. **c)** Summary data of the astrocyte territory area (μm^2^) (Control: 108 astrocytes from 4 mice, Ezr-saCas9: 109 astrocytes from 4 mice). Nested t test (t_191_ = 3.78, ** p = 0.002). **d)** Schematic of experimental workflow of neuron-astrocyte proximity assay (Napa). **e)** Left: representative expression of Napa-n (red) and Napa-a (green) in the CA3 and CA1, respectively. Scale bar: 500 μm. Right: representative image of CA1 astrocytes expressing Napa-a (green) together with HA-saCas9 (magenta), and presynaptic puncta from CA3 projecting neurons (red). Scale bar: 50 μm. **f)** Representative images of CA1 astrocytes from Control and Ezr-saCas9 mice expressing Napa-a (green) in proximity to projections from CA3 neurons expressing Napa-n (red). The FRET ROIs are shown on a color scale that reflects FRET efficiency. Scale bar: 20 μm. **g)** Summary data of the number of FRET signals (ROIs) per CA1 astrocyte (Control: 101 astrocytes from 5 mice, Ezr-saCas9: 141 astrocytes from 5 mice). Nested t test (t_63_ = 3.97, *** p = 0.0002). Data are presented as mean ± SEM.

### Loss of Ezrin reduces astrocyte contacts at synapses

Next, we employed electron microscopy (EM) to closely evaluate the ultrastructure of synapses in the absence of astrocytic Ezrin. Ezr-saCas9 and Control virus were co-injected in the CA1 together with AAV-GfaABC_1_D-GFP (Figure 3a), to confirm sufficient viral expression in samples that were subject to EM analysis (Figure S3a). To determine whether the astroglial synaptic coverage was altered, as already indicated with Napa FRET, we analysed the structural relationship between astrocyte leaflets and synaptic elements. Indeed, we observed that astrocyte-presynapse contact was decreased in Ezr-saCas9 mice compared to controls (Control: 19.7 ± 0.9%, Ezr-Cas9: 16.5 ± 0.6% relative astrocyte-presynaptic contact) (Figure 3b,c). Interestingly, this decrease in astrocyte-synapse contacts was specific for the presynaptic terminal, since astrocyte-postsynapse contact was found to be unaltered (Control: 23.8 ± 1.2%, Ezr-saCas9: 23.9 ± 0.8% relative astrocyte-postsynapse contact) (Figure 3b,d). Furthermore, we found that the leaflet tip, which is in direct opposition of the synaptic cleft, was shorter in Ezr-saCas9 mice (Control: 68.8 ± 4.2, Ezr-saCas9: 50.9 ± 3.3 nm) (Figure 2e,f). Accordingly, we observed an increased distance between the astrocyte leaflet tip and the synaptic cleft in Ezr-saCas9 mice (Control: 75.8 ± 10.2, Ezr-saCas9: 101.1 ± 5.9 nm) (Figure 2g,h). It should be noted that the observed reduction in astrocyte leaflet-synapse interaction in Ezrin-Cas9 mice did not significantly change the number of synapses contacted by an astrocyte (Control: 45.8 ± 3.6%, Ezr-saCas9: 41.7 ± 3.6% of synapses contacted by an astrocyte leaflet) (Figure S3b) nor did it affect the size of the pre-or postsynaptic elements (Control: 1604 ± 25.6, Ezr-saCas9: 1552 ± 21.7 nm (size presynapse); Control: 1378 ± 24. 9, Ezr-saCas9: 1356 ± 20.0 nm (size postsynapse)) (Figure S3c-e). Together these data show that the depletion of Ezrin in astrocytes specifically reduced the interaction of astrocytic leaflets with the presynaptic terminal and the synaptic cleft.

**Figure 3.**
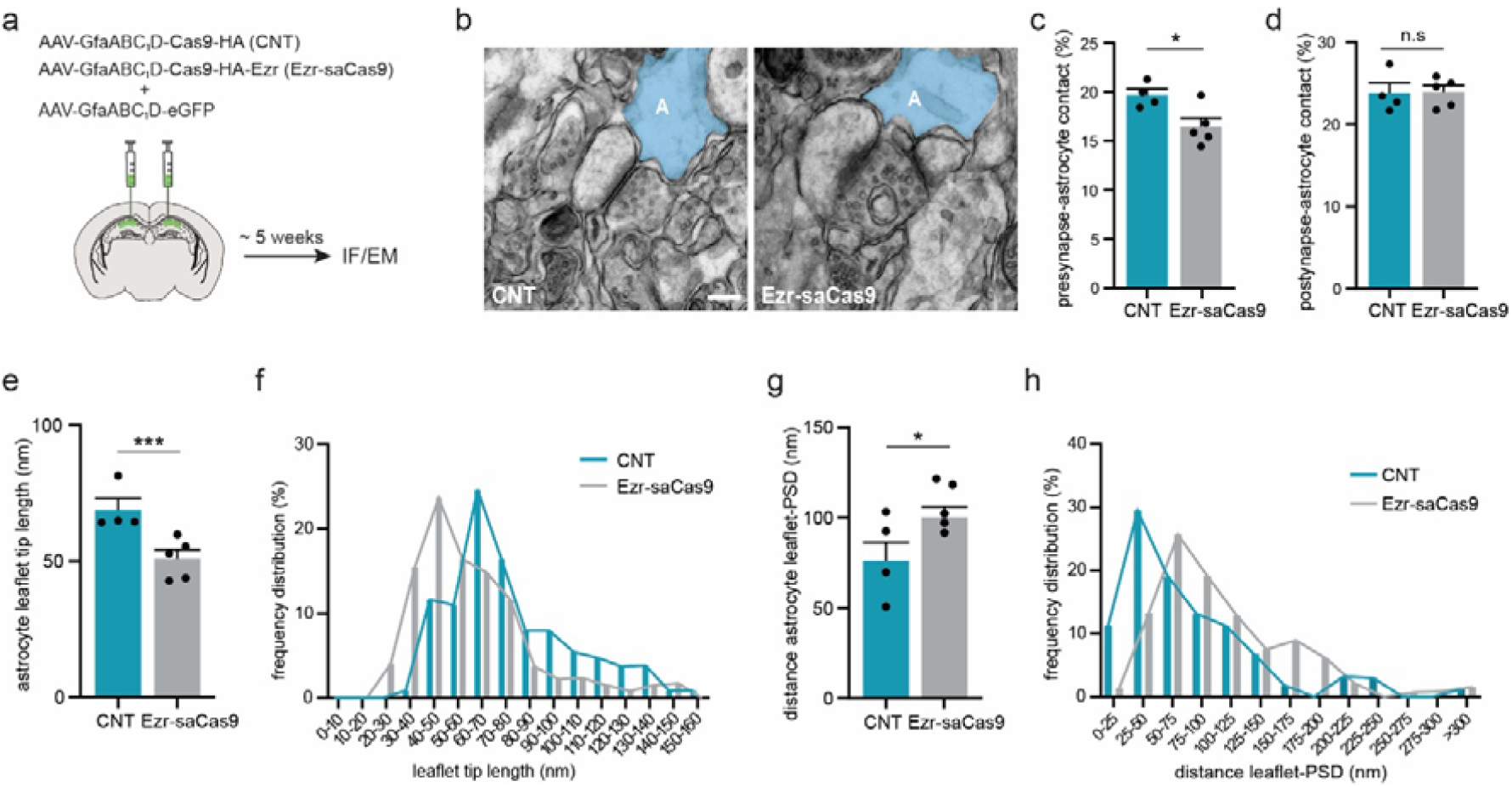
Deletion of Ezrin in hippocampal astrocytes reduced the interaction of astrocytic leaflets with the presynaptic terminal and synaptic cleft. **a)** Schematic of experimental workflow. **b)** Representative images of synapse structure from a Control and Ezr-saCas9 mice. In light blue, an astrocyte leaflet contacting an excitatory synapse. Scale bar: 200 nm. **c)** Summary data of the presynapse and astrocyte contact relative to the presynaptic size from Control and Ezr-saCas9 mice (Control: 157 synapses from 4 mice, Ezr-saCas9: 174 synapses from 4 mice). Nested t test (t_328_ = 2.06, * p = 0.04). **d)** Summary data of the postsynapse and astrocyte contact relative to the postsynapse size from Control and Ezr-saCas9 mice (Control: 193 synapses from 4 mice, Ezr-saCas9: 206 synapses from 4 mice). Nested t test (t_397_ = 0.35, p = 0.72), n.s = not significant. **e)** Summary data of astrocyte leaflet tip length from Control and Ezr-saCas9 mice (also see Figure S3f) (Control: 101 astrocytes from 4 mice, Ezr-saCas9: 116 astrocytes from 4 mice). Nested t test (t_157_ = 3.67, *** p = 3.67). **f)** Frequency distribution plot from data presented in **e**. Note that Ezr-saCas9 mice show a clear shift towards a smaller astrocyte leaflet tip. **g)** Summary data of the distance between the astrocyte leaflet tip and the post-synaptic density (PSD) from Control and Ezr-saCas9 mice (also see Figure S3g) (Control: 96 astrocytes from 4 mice, Ezr-saCas9: 110 astrocytes from 4 mice). Nested t test (t_148_ = 2.39, * p = 0.01). **h)** Frequency distribution plot from data presented in **g**. Note that Ezr-saCas9 mice show a shift towards larger distances. Data are presented as mean ± SEM.

### Disrupted astroglial coverage decreases synaptic glutamate levels

To examine whether Ezr-saCas9 mediated manipulation of astrocyte leaflet structure has consequences for synaptic function, we analyzed excitatory synaptic transmission 5 weeks after viral delivery (Figure 4a). Whole-cell patch-clamp recordings from hippocampal pyramidal neurons revealed no change in sEPSC amplitude (Control: 171.4 ± 21.7, Ezr-saCas9: 151.1 ± 16.2 pA) and frequency (Control: 15.8 ± 0.7, Ezr-saCas9: 14.9 ± 0.6 s), suggesting that basal synaptic transmission was unaffected in Ezr-saCas9 mice (Figure 4b-d). Next, we measured evoked AMPAR-mediated EPSCs and synaptic glutamate levels in Ezr-saCas9 mice. For this, CA3-CA1 Schaffer Collaterals were stimulated with or without γ-DGG, a low-affinity competitive antagonist of AMPARs, at a non-saturating concentration (1 mM) (Liu et al., 1999). At this concentration, effectiveness of the drug depends on synaptic glutamate levels. Evoked EPSC amplitude (Control: 171.4 ± 21.7, Ezr-saCas9: 151.1 ± 16.2 pA) and decay kinetics (Control: 8.7 ± 0.40 and Ezr-saCas9: 9.0 ± 0.5 ms) were not changed prior to drug application (Figure S4a-c). However, we found that γ-DGG inhibition of evoked AMPAR EPSCs was more pronounced in pyramidal cells from Ezr-saCas9 mice than from controls (Control: 45.3 ± 5.0%, Ezr-saCas9: 59.1 ± 4.2% of inhibition) (Figure 4e-h). Together, this shows that basal strength of synaptic transmission was unaffected in Ezr-saCas9 mice, but that evoked synaptic glutamate levels were lower in Ezr-saCas9 mice compared to controls. It has been reported that astrocyte process withdrawal from synapses in the supraoptic nucleus causes a local build-up of glutamate and subsequent activation of presynaptic metabotropic receptors, leading to a reduced presynaptic release probability (Oliet, 2001). To determine whether a reduction in presynaptic release probability underlies the observed y-DGG effect in Ezr-saCas9 mice, we examined the paired-pulse ratio (PPR) and found no significant PPR differences between Ezrin-saCas9 mice and controls (Figure 4i,j). Taken together, these data show that Ezr-saCas9 mice have reduced evoked synaptic glutamate levels, which is unlikely due to a presynaptic release defect.

**Figure 4.**
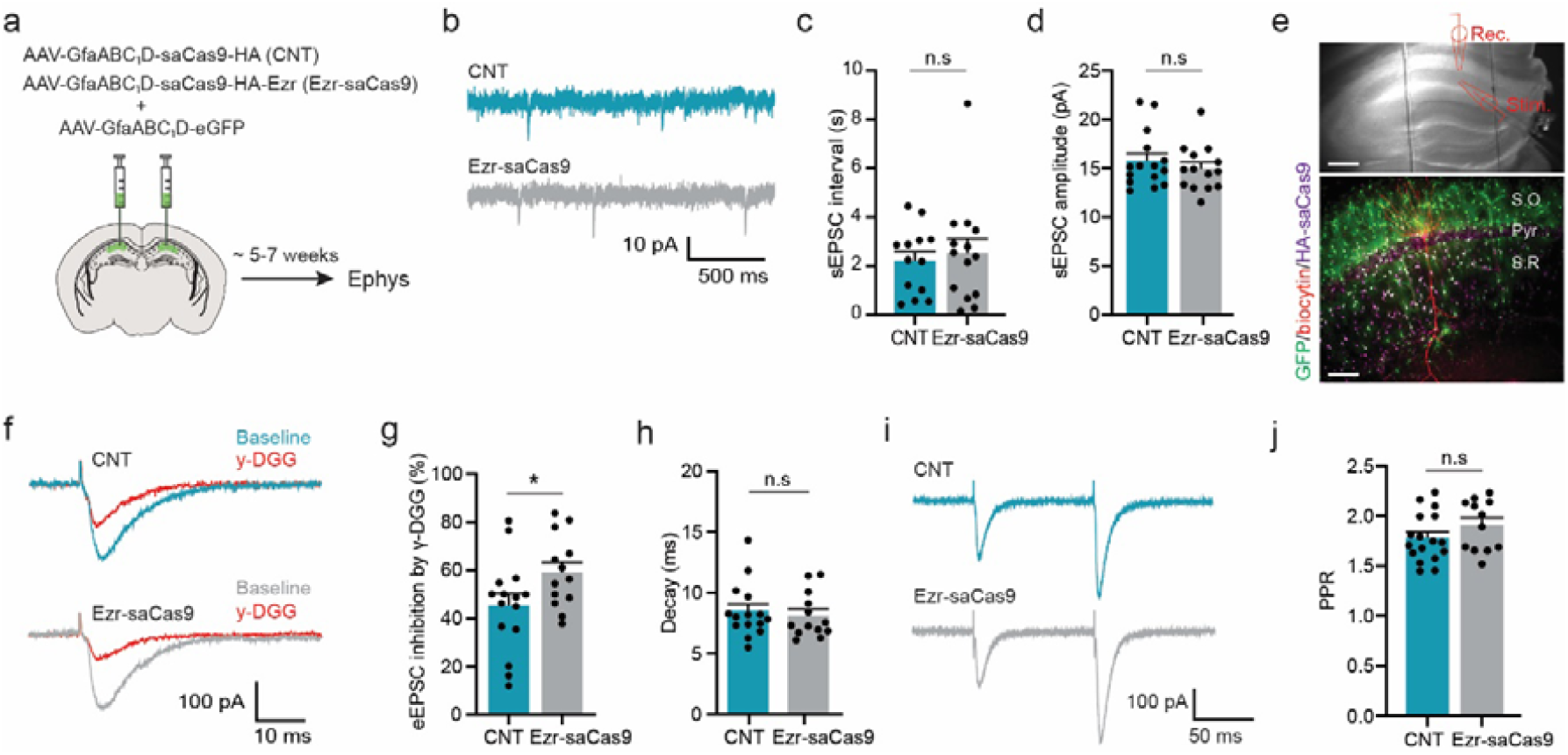
Reduced astroglial coverage due to loss of Ezrin does not affect basal synaptic transmission but it decreases evoked synaptic glutamate levels. **a)** Schematic of experimental workflow. **b)** Representative traces of spontaneous AMPAR EPSCs from pyramidal neurons from Control and Ezr-saCas9 mice. **c**,**d)** Summary data of (**c**) interval and (**d**) amplitude of sEPSCs (Control: 13-14 cells from 7 mice, Ezr-saCas9: 14-15 cells from 7 mice). Unpaired t test: (interval) t_25_ = 0.45, p = 0.65; (amplitude) t_27_ = 0.81, p = 0.42; n.s = not significant. **e)** Upper: representative image of the dorsal hippocampus with an adjacently placed glass pipette (rec.). The stimulation glass pipette (stim.) was placed in the *Stratum Radiatum*. Scale bar: 200 μm. Bottom: representative image of a patched neuron filled with biocytin (red) surrounded by astrocytes expressing GFP and HA-saCas9. Scale bar: 100 μm. **f)** Representative traces of evoked AMPAR EPSCs (eEPSCs) from pyramidal neurons from Control and Ezr-saCas9 mice with and without _γ_-DGG (1 mM). **j**,**h)** Summary data of the (**j**) percentage of eEPSC amplitude inhibited by _γ_-DGG and (**h**) decay kinetics in presence of _γ_-DGG (Control: 15 cells from 6 mice, Ezr-saCas9: 13 cells from 6 mice). Unpaired t test: (amplitude) t_26_ = 2.087, * p = 0.04; (decay) t_26_ = 0.46, p = 0.64; n.s = not significant. **i)** Representative traces of paired pulse ratio (PPR) from Control and Ezr-saCas9 mice. **j**) Summary data of PPR from Control and Ezr-saCas9 mice (Control: 17 cells from 7 mice, Ezr-saCas9: 12 cells from 7 mice). Unpaired t test: t_27_ = 1.38, p = 0.17. Data are presented as mean ± SEM.

### Increased glutamate spill over and NMDA receptor activation upon deletion of astrocytic Ezrin

To investigate how the reduced synaptic glutamate levels following evoked stimulation relate to increased glutamate spillover, as a consequence of reduced astrocyte leaflet-synapse interaction, the temporal dynamics of extrasynaptic glutamate were investigated. For this, the glutamate sensor iGluSnFR (Marvin et al., 2013) was expressed on CA1 astrocytes, together with either Ezr-saCas9 or control virus (Figure 5a). Two-photon imaging of iGluSnFR signals allowed study of the time course of extrasynaptic glutamate following synaptic stimulation. Synaptic activity was evoked by focal electrical stimulation (10 pulses at 50 Hz) in the CA1. The stimulation electrode was placed in proximity of an iGluSNFR expressing astrocyte and glutamate transients were measured within a circular region of interest (ROI) on the astrocyte domain (Figure 5b). We did not observe significant changes in the magnitude of iGluSnFR signal between groups (Control: 1.2 ± 0.01, Ezr-saCas9: 1.3 ± 0.02 ΔF/F) (Figure S5a,b). To examine glutamate temporal dynamics, we fitted the decay of the averaged evoked glutamate transients (6 sweeps) with a single exponential (Figure 5c) and found that the glutamate transients in Ezr-saCas9 mice displayed an increased decay time following trains of 50 Hz stimulation (Control: 71.3 ± 6.3, Ezr-saCas9: 91.2 ± 7.0 ms) (Figure 5d,e). The decay of the iGluSnFR transients was not influenced by the amplitude of the response (Figure S5c,d). Together, these data suggest increased dwelling of glutamate in the extrasynaptic space which is in line with the observed decrease in astroglial coverage of excitatory synapses in Ezr-saCas9 mice.

**Figure 5.**
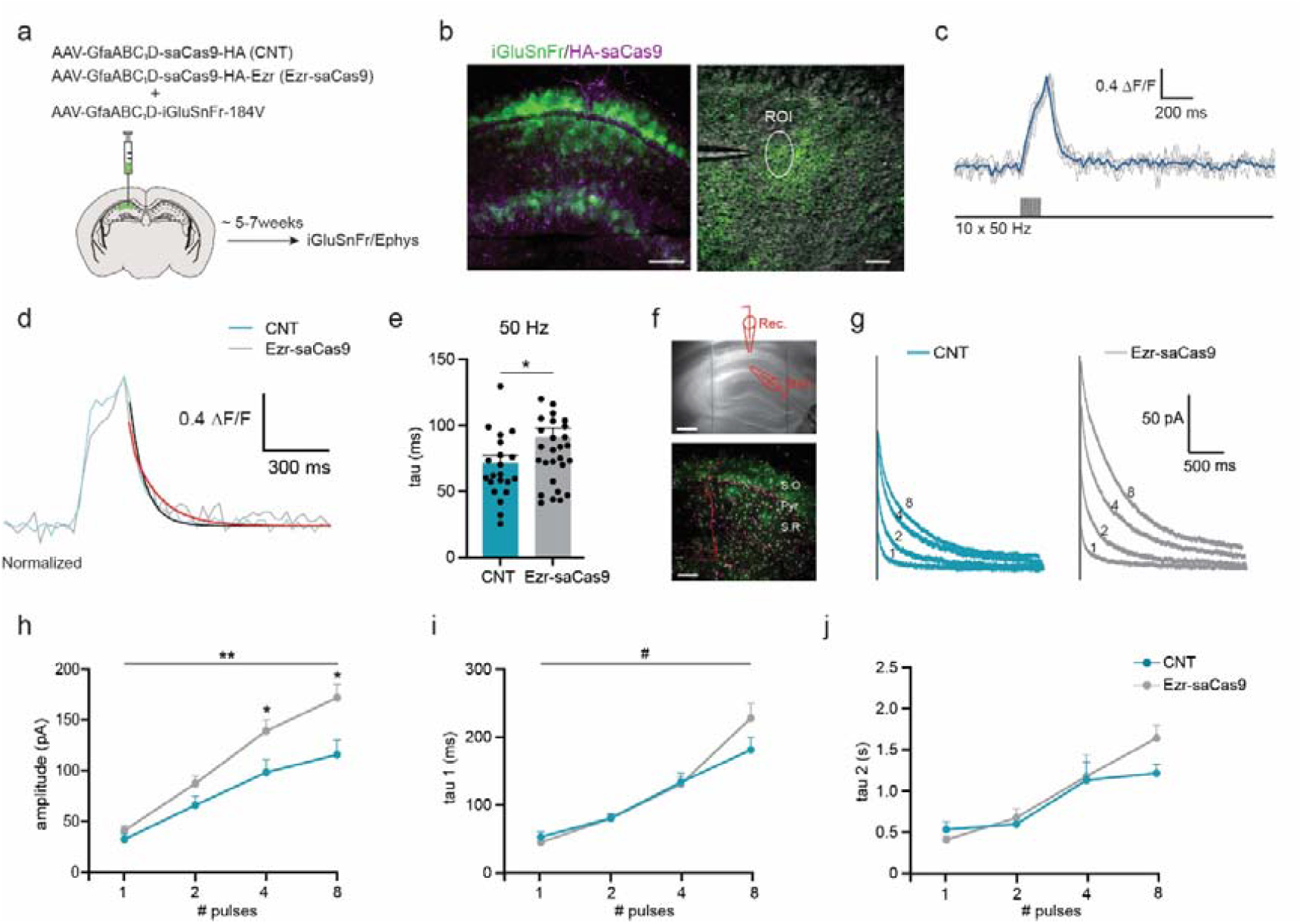
Decreased astroglial synaptic coverage boosts glutamate spillover and NMDAR activation. **a)** Schematic of experimental workflow. **b)** Left: representative images of CA1 astrocytes expressing AAV-GfaABC_1_D::iGluSnFR and AAV-GfaABC_1_D::Cas9-HA-Ezr. Scale bar: 200 μm. Right: stimulation glass pipette was placed adjacently to an astrocyte expressing iGluSnFR and transients were measured and quantified from the region of interest (ROI). Scale bar: 20 μm. **c**,**d)** Upon synaptic stimulation (10 × 50 Hz), a robust increase in iGluSnFR signal was detected. **c)** Thick blue line represents the average of 6 responses. **d)** Thick black (Control) and red (Ezr-saCas9) lines represent the single-exponential fit of the decay. **e)** Summary data of the decay kinetics (Control: 22 astrocytes from 7 mice, Ezr-saCas9: 31 astrocytes from 7 mice) following 50 Hz stimulation. Unpaired t test: t_51_ = 2.06, * p = 0.02). **f)** Upper: representative image of the dorsal hippocampus with an adjacently placed glass pipette (rec.). The stimulation glass pipette (stim.) was placed in the *Stratum Radiatum*. Scale bar: 200 μm. Bottom: representative image of a patched neuron filled with biocytin (red) surrounded by astrocytes expressing AAV-GfaABC_1_D::GFP and AAV-GfaABC_1_D::saCas9-HA-Ezr. Scale bar: 100 μm. **g)** Representative traces of the NMDAR EPSCs upon incremental short burst stimulation (1, 2, 4, 8) for Control and Ezr-saCas9 mice. **h-j)** Summary data of the (**h**) amplitude and (**i**,**j**) kinetics of the NMDAR-mediated currents upon incremental short burst stimulation (1, 2, 4, 8) (Control: 17 cells from 7 mice, Ezr-saCas9: 27 cells from 9 mice). Amplitude (**h**): repeated measures two-way ANOVA, interaction effect Pulses x Genotype; F_(1.21,15.38)_ = 8.37, ** p = 0.0082. Post-hoc Bonferroni test Control vs Ezr-saCas9: 1 pulse p = 0.41, 2 pulses p = 0.15, 4 pulses * p = 0.04, 8 pulses * p = 0.03. Tau 1 (**i**): repeated measures two-way ANOVA, interaction effect Pulses x Genotype; F_(1.76,22.29)_ = 3.04, # p = 0.07. Post-hoc Bonferroni test Control vs Ezr-saCas9: 1 pulse p = 0.99, 2 pulses p = 0.99, 4 pulses p = 0.99, 8 pulses p = 0.32. Tau 2 (**j**): repeated measures two-way ANOVA, interaction effect Pulses x Genotype; F_(1.7,23.84)_ = 1.11, p = 0.33. Post-hoc Bonferroni test control vs Ezr-saCas9: 1 pulse p = 0.99, 2 pulses p = 0.99, 4 pulses p = 0.99, 8 pulses p = 0.31. Data are presented as mean ± SEM.

As astrocyte leaflets contain the high-affinity glutamate transporter GLT-1, we next tested whether the lack of Ezrin might slow glutamate clearance. We found that partial blockade of glutamate transporters (GluTs) with threo-beta-Benzyloxyaspartate (DL-TBOA) (10 μM) significantly similarly increased the decay kinetics of glutamate transients in both groups (Control: 134.2 ± 10.1, Ezr-saCas9: 145.6 ± 10.8 ms (post-TBOA)) (Figure S5e,f). Thus, these data suggest that even though astrocyte leaflets are further away from the synaptic cleft in Ezr-saCas9 mice, the uptake of glutamate remains unchanged.

Perisynaptic glutamate has been shown to increase the activation of extrasynaptic NMDA receptors (Kullmann and Asztely, 1998). We therefore stimulated CA1 pyramidal neurons with incremental short bursts (1-2-4-8 stimuli) of high frequency stimulation (100 Hz) to increase glutamate spillover (Lozovaya et al., 2004), and measured NMDAR-mediated EPSCs (Figure 5a,f,g). Ezr-saCas9 mice showed a remarkable increase in NMDAR-mediated EPSCs amplitude (Control: (1 pulse) 32.6 ± 5.5; (2 pulses) 66.1 ± 9.1; (4 pulses) 98.4 ± 12.3; (8 pulses) 115.8 ± 14.3 pA, Ezr-saCas9: (1 pulse) 41.2 ± 3.3; (2 pulses) 87.4 ± 7.3; (4 pulses) 139.3 ± 11.1; (8 pulses) 171.9 ± 13.1 pA) (Figure 5g,h) accompanied with a slightly slower decay (Control: (1 pulse) 58.1 ± 5.3; (2 pulses) 89.6 ± 4.3; (4 pulses) 146.0 ± 14.3; (8 pulses) 199.5 ± 20.8 ms, Ezr-saCas9: (1 pulse) 50.4 ± 4.1, (2 pulses) 90.4 ± 8.2, (4 pulses) 146.2 ± 12.3, (8 pulses) 257.9 ± 23.7 ms) (Figure 5i,j). For both amplitude and decay, the difference in response between Ezr-saCas9 and controls became larger with longer bursts of synaptic stimulation, which enhances glutamate spillover from the synaptic cleft. These results are consistent with the interpretation that reduced astrocyte leaflet-synapse interaction leads to enhanced synaptic cooperation (via activation of extrasynaptic NMDARs and/or NMDARs in neighboring synapses) (Arnth-Jensen et al., 2002; Henneberger et al., 2020). Together, we show that Ezrin-mediated astrocytic leaflet manipulation increases the time course of glutamate in the extrasynaptic space and leads to increased NMDAR activation.

### Reduced astroglial coverage enhances recent contextual fear memory retrieval and increases neuronal activity

Previous research has shown that leaflet invasion towards the synaptic cleft impaired recent contextual fear memory recall (Pannasch et al., 2014). Hence, we explored whether reduced astroglial coverage altered contextual fear memory expression. Mice were fear conditioned 5 weeks after viral delivery, and memory expression was measured at recent and remote time-points after learning (Figure 6a). Interestingly, Ezr-saCas9 mice exhibited enhanced fear expression when compared to control mice 5 days, but not 28 days after contextual fear conditioning (CFC) (Control: 54.7 ± 3.2%, Ezr-saCas9: 70.8 ± 4.5% of freezing 5 days after training; Control: 57.1 ± 10.3%, Ezr-saCas9: 65.4 ± 7.3% of freezing 28 days after training) (Figure 6b and Figure S6a). No differences were found between controls and Ezr-saCas9 mice in the open field (exploratory behavior and the time spent at the center of the arena), indicating that the observed phenotype was not driven by anxiety or locomotor deficits (Control: 31.1 ± 2.1, Ezr-saCas9: 27.4 ± 1.8 m; Control: 17.5 ± 2.4%, Ezr-saCas9: 20.1 ± 2.0% time spent in the center) (Figure S6b-d). Furthermore, no effect on freezing levels were detected in an independent cohort of mice that was trained in the conditioning box but placed in a novel context 5 days later (Control: 18.85 ± 3.77%, Ezr-saCas9: 13.24 ± 4.43% of freezing in the novel context) (Figure 6a,c), demonstrating that the enhanced fear memory observed in Ezr-saCa9 mice was indeed context-specific and time-dependent.

**Figure 6.**
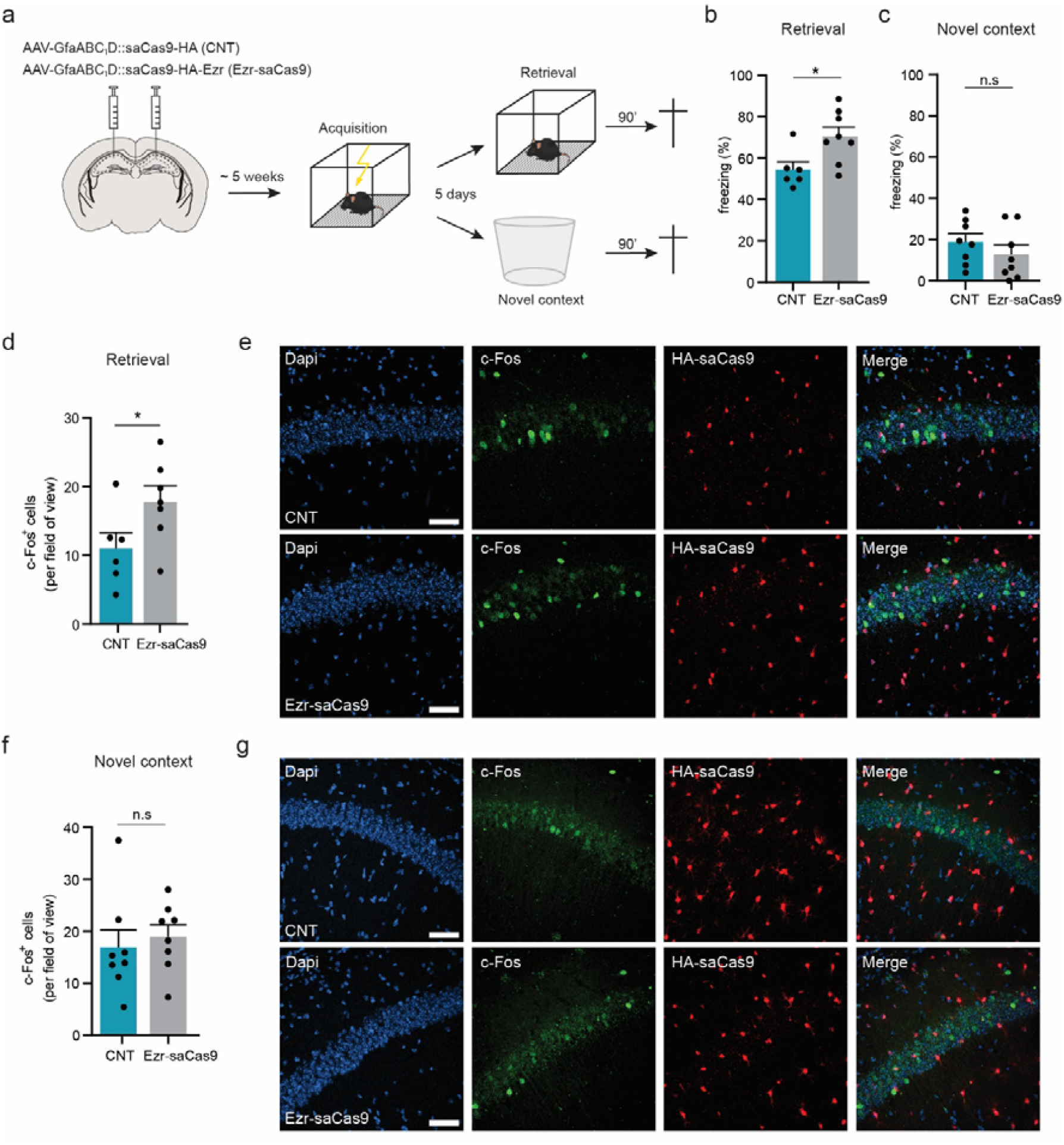
Enhanced recent contextual fear memory and c-Fos^+^ cells in Ezr-saCas9 mice. **a)** Schematics of experimental workflow. **b)** Freezing levels during retrieval (Control: 6 mice and Ezr-saCas9: 8 mice). Unpaired t test: t_12_ = 2.63, * p = 0.02. **c)** Freezing levels during novel context (Control and Ezr-saCas9: 8 mice). Unpaired t test: t_14_ = 0.96, p = 0.35). **d)** Summary data of c-Fos^+^ cells in the CA1 pyramidal layer during recent retrieval (Control: 8 slices per mouse from 6 mice, Ezr-saCas9: 8 slices per mouse from 7 mice). Unpaired t test: t_11_ = 2.27, * p = 0.04. **e)** Representative examples of activated neurons (c-Fos^+^ cells) in the CA1 during recent retrieval. **f)** Summary data of c-Fos^+^ cells in the CA1 pyramidal layer during novel context exposure (Control: 8 slices per mouse from 7 mice, Ezr-saCas9: 8 slices per mouse from 8 mice). Unpaired t test: t_14_ = 0.52, p = 0.6. **g)** Representative examples of activated neurons (c-Fos^+^ cells in the CA1 during novel context. Scale bar: 50 μm. Data are presented as mean ± SEM.

Next, we tested whether memory enhancement was accompanied by increased neuronal activity in a context-dependent manner. For this, we measured c-Fos levels (a marker for neuronal activation (Cruz et al., 2015)) in CA1 neurons of mice that underwent retrieval in the conditioning context, those that were placed in novel context during retrieval, and naïve home-cage (HC) controls. In line with the enhanced freezing levels observed, Ezrin-saCas9 mice had a greater number of c-Fos^+^ neurons 90 minutes after recent retrieval when compared to controls (Control: 10.7 ± 2.2, Ezr-saCas9: 18 ± 2.3 c-Fos^+^ neurons in the CA1) (Figure 6d,e). Notably, this increased neuronal activation was context- and memory-specific, as no differences were observed between groups after retrieval in a novel context (Control: 16.8 ± 2.3, Ezr-saCas9: 18.9 ± 3.4 c-Fos^+^ neurons in the CA1) (Figure 6f,g), nor in naïve HC controls (Control: 2.3 ± 0.2, Ezr-saCas9: 3.4 ± 0.7 c-Fos^+^ neurons in the CA1) (Figure S6e,f). Together, these data demonstrate that Ezrin-dependent astrocyte leaflet-synapse interactions influence recent memory retrieval and gate neuronal activation in a context-dependent manner.

## Discussion

In the present study we used genetic and imaging tools to manipulate astrocyte leaflets *in vivo*, and discovered that these thin astrocytic structures that contact synapses are critical for recent memory expression. Specifically, we identified Ezrin as a protein required for intact astrocyte morphology and proximity to synapses in the adult hippocampus. Furthermore, by interfering with astrocyte leaflet structure, we show that they are necessary in the gating of glutamate spillover and the expression of recent contextual memory.

Recent studies have demonstrated a crucial role for astrocyte-neuron signalling in shaping neuronal circuits and behaviour (Adamsky et al., 2018; Gao et al., 2016; Kol et al., 2020; Martin-Fernandez et al., 2017; Mederos et al., 2021; Nagai et al., 2019; Papouin et al., 2017; Robin et al., 2018; Suzuki et al., 2011). Much of this work is based on the notion that astrocytes processes are located in close proximity to synapses. However, how this proximity affects cognitive performance has remained largely unexplored (Pannasch et al., 2014; Zhou et al., 2019). We employed the CRISPR-Cas9 gene editing to manipulate astrocyte processes by targeting astrocytic Ezrin in the adult hippocampal CA1 region. We found the viral Ezr-saCas9 vector to have a high penetrance and nearly complete specificity, resulting in ∽73% reduction of Ezrin at both mRNA and protein level five weeks after viral transduction. Thus, the current approach enabled us to optimally and selectively reduce Ezrin expression in a large population of hippocampal astrocytes and to bypass any developmental role of Ezrin. It should be noted, however, that the Ezr-saCas9 vector did not lead to a complete absence of astrocytic Ezrin, therefore, some of the effects reported in our study are likely to be an underestimation.

Ezrin was previously found to be specifically localized to astrocyte leaflets (Derouiche and Frotscher, 2001) and it was demonstrated to be required for filopodia formation in cultured astrocytes (Lavialle et al., 2011). Our results support previous findings in the developing somatosensory cortex (Zhou et al., 2019) and suprachiasmatic nucleus (Lavialle et al., 2011) where Ezrin expression correlated with astrocyte morphogenesis and leaflet formation (Zhou et al., 2019). Here, we built on these data to demonstrate that Ezrin is required for astrocyte leaflets structure and thereby synaptic function. We show that the reduction of Ezrin in adult hippocampal astrocytes compromises astrocyte morphogenesis and reduces astrocyte leaflet-synapse contact. We provide new evidence that decreased astrocytic Ezrin expression in adult mice leads to reduced astroglial coverage of hippocampal excitatory synapses, predominantly affecting both the astrocyte-presynapse contact and the interaction of the leaflet tip with the synaptic cleft. It is thought that astrocyte leaflets form stable contacts with dendritic spines (Arizono et al., 2020) to support spine morphogenesis (Murai et al., 2003) and synaptic function (Filosa et al., 2009), and therefore, astrocytes that exhibit shorter leaflets may be most likely to disengage from the presynapse rather than the postsynapse. These observations imply that astrocytic Ezrin could link membrane-bound proteins with actin filaments, and may thereby be essential in the formation and motility of the fine glial processes upon specific stimuli (Derouiche and Geiger, 2019). This mechanism has been suggested for other actin binding proteins, such as Cofilin-1, that regulates leaflet shrinkage through NKCC1 (Na^+^, K^+^, 2Cl^-^ cotransporter) activation (Henneberger et al., 2020), and Profilin-1, that regulates leaflet outgrowth upon Ca^2+^ elevation (Molotkov et al., 2013).

Previous reports on astrocyte-synapse interaction performed in acute brain slices revealed that astrocyte leaflets undergo structural rearrangements upon synaptic activation as a mechanism to modulate local synaptic transmission and plasticity (Bernardinelli et al., 2014; Henneberger et al., 2020; Lushnikova et al., 2009; Perez-Alvarez et al., 2014). Since our manipulation affected the spatial organization of leaflets at synapses, we examined excitatory synaptic transmission. First, we found that the lack of astrocytic Ezrin had no effect on basal synaptic transmission. In line with this, reduction of astrocyte territory and astrocyte-synapse interaction by deletion of the astrocyte transcription factor NFIAA, did also not influence hippocampal basal synaptic function (Huang et al., 2020). In fact, Ezrin-dependent astrocyte leaflet-synapse interaction becomes only apparent in response to strong activation of local synapses; e.g. incremental burst stimulation that leads to NMDARs activation (Figure 5) or learning-induced synaptic activation (Figure 6). Thus, the increased NMDAR mediated responses in Ezrin-saCas9 mice are likely a consequence of increased glutamate spillover into the extracellular space during repetitive synaptic stimulation. This is consistent with previous findings in hippocampal acute slices, where LTP-induced leaflet withdrawal enhanced the activation of extrasynaptic NMDARs and facilitated NMDAR-mediated cross-talk among neighbouring synapses (Henneberger et al., 2020). Additionally, we show that the increased glutamate spillover and NMDAR activation in the Ezr-saCas9 mice likely reflects an increased diffusion in the extracellular space (Piet et al., 2004) as glutamate clearance does not seem to be impaired. Taken together, our data reveals that a decrease in astrocytic synaptic coverage does not affect basal synaptic transmission, but is sufficient to boost glutamate spillover and extrasynaptic NMDAR activation under conditions of robust neuronal activation.

Recent work has demonstrated the importance of astrocyte-neuronal communication for cognitive function in both healthy (Adamsky et al., 2018; Gao et al., 2016; Gibbs et al., 2006; Kol et al., 2020; Martin-Fernandez et al., 2017; Mederos et al., 2021; Nagai et al., 2019; Santello et al., 2019) and diseased (Lioy et al., 2011; Orr et al., 2015; Reichenbach et al., 2018; Takahashi et al., 2015) brains. Yet, it remained poorly understood whether changes in proximity of the astrocyte leaflet to the synapse affects learning and memory. Previous studies monitoring leaflet-synapse interaction upon neurotransmitter release or synaptic plasticity were time restricted (from minutes to two hours) (Bernardinelli et al., 2014; Henneberger et al., 2020; Lushnikova et al., 2009; Perez-Alvarez et al., 2014) or solely investigated in acute brain slices (Lushnikova et al., 2009; Perez-Alvarez et al., 2014). Nonetheless, Pannasch et al (2014) showed that deletion of Cx30 causes astroglial invasion of the synaptic cleft and subsequently reduces synaptic strength and impairs contextual fear memory retrieval (Pannasch et al., 2014). It should be noted that increased leaflet invasion of the synaptic cleft has so far not been observed under physiological conditions. Here, conversely, we demonstrate that when the astrocyte leaflet is further away from the synaptic cleft, mice exhibit increased glutamate spillover and enhanced memory recall. Interestingly, we observe that the enhanced retrieval is time and context specific, which is in line with previous studies demonstrating the importance of the hippocampus in the synaptic consolidation of recent context memories (Frankland and Bontempi, 2005; Liu et al., 2012). Furthermore, Ezr-saCas9 mice showed a context-specific increase in neuronal activity, suggesting that more neurons were activated at the time of retrieval, supporting the notion that astrocytes show a tailored response to the activity of their surrounding neurons (Adamsky et al., 2018). Thus, our data reveal that the proximity of the astrocyte leaflets to the synaptic cleft determines the extent of memory expression and the activation of its supporting neurons. We propose that astrocyte leaflet-synapse interaction plays a substantial role in synaptic consolidation, by supporting learning-induced remodelling of pre-existing neuronal connections and the formation of new ones (Rao-Ruiz et al., 2021). Accordingly, learning-induced increased in glutamate spillover, spillover, due to reduced astroglial synaptic coverage, might promote the engagement of synapses that are in close proximity, and consequently, enhance synaptic connectivity that facilitates memory retrieval (Govindarajan et al., 2006). Whether learning is dependent on structural rearrangement of astrocyte leaflets is an intriguing and important question that remains to be addressed.

In summary, to our knowledge, this study is the first to show that a selective and physiologically relevant manipulation of astrocyte leaflets structure leads to enhanced memory retrieval in adult mice. Our data support the proposition that the proximity of astrocyte processes with synapses in the adult brain is instrumental in the plasticity mechanisms that underlie memory performance.

## Acknowledgments

The authors would like to thank Yvonne Gouwenberg and Robbert Zalm (VU University, Amsterdam, The Netherlands) for AAV vector construct preparation, Rolinka van der Loo (VU University, Amsterdam, The Netherlands) for helping with animal perfusions, J Christopher Octeau for all the technical support during Napa experiments, and Priyanka Rao-Ruiz (VU University, Amsterdam, The Netherlands) for the critical revision of the manuscript.

## Author contributions

A.B.S performed stereotaxic surgeries, histology, confocal imaging, NAPA, two-photon glutamate imaging and all behavioural experiments. M. S. J. K performed electron microscopy experiments. T.S.H performed all electrophysiological experiments. B.K co-supervised the Napa experiments. R.M co-supervised electrophysiological and two-photon experiments. A.N built the two-photon microscope. M.H.G.V and A.B.Smit conceived and supervised all aspects of the project and secured funding. A.B.S and M.H.G.V wrote the manuscript with input from all the other authors.

## Methods

### Animals

Wild-type, male C57BL/6J mice were 8-10 weeks old at the start of experiments and were individually housed on a 12 h light/dark cycle with ad libitum access to food and water.

Behavioral experiments were performed during the light phase. All experimental procedures were approved by The Netherlands central committee for animal experiments (CCD) and the animal ethical care committee (DEC) of the Vrije Universiteit Amsterdam (AVD1120020174287). Mice were randomly assigned to experimental groups.

### Constructs

The pAAV-GfaABC_1_D::Cas9-HA-Ezr was generated by first replacing the CMV promoter from pX601-AAV-CMV::NLS-saCas9-NLS-3xHA-bGHpA;Bsal-sgRNA (Addgene plasmid #61591) with the GFAP promoter (GfaABC_1_D). Next, we inserted the designed sgRNA to target exon-1 of Ezrin (TGGCTGGTTGGTGGCTCTGCGTGGGT) (Genscript: NM_001271663.1_T3). Finally, we cloned the modified plasmid to an AAV2/5 vector. The control virus, pAAV-GfaABC_1_D::Cas9-HA, was generated following the same procedure but it lacks the sequence to target Ezrin.

### AAV vectors and stereotaxic micro-injections

AAV-GfaABC_1_D::Cas9-HA-Ezr (titer (GC/ml): 2.6 × 10^10^, 0.7 μl per site), AAV-GfaABC_1_D::Cas9-HA (titer (GC/ml): 7.64 × 10^10^, 0.7 μl per site), AAV-GfaABC_1_D::eGFP (titer (GC/ml): 8.3 × 10^12^, diluted 1:20 with sterile PBS, 0.5 μl per site), AAV-GfaABC_1_D::SF-iGluSnFR.A184V (plasmid from Addgene #106193) (titer (GC/ml): 5.53 × 10^10^, 0.7 μl per site), pZac2.1-GfaABC_1_D::NAPA-A SV40 (gift from Baljit Khakh, Addgene #92281) (titer (GC/ml): 1.36 × 10^11^, diluted 1:5 with sterile PBS, 0.7 μl left hemisphere), were packaged as serotype 2/5 virus. pZac2.1-hSynapsin1::NAPA-N SV40 (gift from Baljit Khakh, Addgene #92282) (titer: 2 × 10^11^, diluted 1:2 with sterile PBS, 0.3 μl left hemisphere) was packaged as serotype 2/1 virus. For stereotaxic micro-injections, mice received 0.1 mg per kg Temgesic (RB Pharmaceuticals, UK) 30 minutes prior to start, and were then anesthetized with isoflurane and mounted onto a stereotactic frame. Lidocaine (2%, Sigma-Aldrich Chemie N.V, The Netherlands) was topically applied to the skull to provide local analgesia. The AAV vectors were infused bilaterally, except AAV2/5-GfaABC_1_D::NAPA-A SV40 that was infused unilaterally, in the CA1 region of the hippocampus (AP: -1.75 mm, DV: -1.7, ML: ± 1.3; relative to Bregma), and AAV2/1-hSynapsin1::NAPA-N SV40 was infused unilaterally in the CA3 region of the hippocampus (AP: -1.8 mm, DV: -2.55, ML: ± 2.4; relative to Bregma). The desired volume was infused using a microinjection glass needle at a rate of 0.1 μl / min. Following infusion, the needle was left in place for 5 minutes to allow virus diffusion and prevent backflow. After surgery, mice remained in their home-cage for 4-5 weeks until the start of experiments.

### Electron microscopy

Mice aged 8-10 weeks were injected bilaterally with 0.7 μl of AAV2/5-GfaABC_1_D::Cas9-HA-Ezr or AAV2/5-GfaABC_1_D::Cas9-HA together with AAV2/5-GfaABC_1_D::eGFP (3:1). Mice were transcardially perfused with 4% paraformaldehyde (PFA) in 0.1M PBS (pH 7.4). Brains were dissected and post-fixated in 4% PFA for 24 h followed by cryo-preservation in 30% sucrose and stored at -80 °C until further processing. 50 μm thin coronal slices of the hippocampus were made on a cryostat and kept in PBS for a maximum of 24 h. Free-floating sections were contrasted with 1% osmium tetroxide and 1% ruthenium. Consequently, the sections were dehydrated by increasing ethanol concentrations (30, 50, 70, 90, 96, 100%) and propylene oxide followed by embedding in epoxy resin and incubation for 72 h at 65 °C. An ultra-microtome (Reichert-Jung, Ultracut E) was used to generate 90 nm thin slices of the hippocampus CA1 region. Finally, the slices were post-contrasted with uranyl acetate and lead citrate in an ultra stainer (LEICA EM AC20).

A JEOL1010 transmission electron microscope was used to acquire digital images of the hippocampus CA1 *stratum radiatum* at 50.000 x magnification. Per animal, a total of 100 randomly selected images were captured and 130 morphologically intact synapses were identified for analysis, from which the presynapse and postsynapse perimeter were measured. When a synapse was recognized, the contact between presynapse and/or postsynapse and astrocyte leaflet was measured. When the astrocyte leaflet was contacting the synaptic cleft, the following parameters were quantified: the distance between astrocyte leaflet and postsynaptic density (PSD) and the astrocyte leaflet tip length. All measurements were performed in ImageJ.

### Neuron-astrocyte proximity assay (NAPA) and FRET analyses

FRET from acute slices was examined 4-6 weeks following AAV injections *in vivo*. We measured FRET by sensitized emission (SE-FRET) using PixFRET, and ImageJ Plug-in. Image processing was carried out as described previously (Badia□Soteras et al., 2020).

Images were acquired using a Nikon confocal microscope with a 40 x, 0.85 numerical aperture water immersion objective. The excitation wavelength used for Napa-a (GFP) was 488 nm and fluorescence was detected at 525/550-nm (I_donor_), while for Napa-n (mCherry) the excitation wavelength was 560 nm, and fluorescence was detected at 595/650 nm (I_acceptor_). To detect SE-FRET, we measured an additional component, which is the total signal emitted at 595/650 nm when excited at 488 nm (I_FRET_). All images were collected with a scan zoom of 3, a frame size of 512 × 512 pixels, a pinhole diameter of 46.5 μm, and 0.5 frames per second. Laser power, gain and offset were optimized to achieve the maximal signal-to-noise ratio and were kept constant for all the samples, including bleed through (Napa-n only) and cross-talk (Napa-a only) controls. We quantified FRET signals in the CA1 *stratum radiatum*. FRET ROIs with a donor/acceptor ratio <0.1 and >10 were discarded, as they lead to an under-and over-estimation of FRET efficiency(Octeau et al., 2018). We quantified the number of FRET ROIs per astrocyte domain.

### Slice preparation and electrophysiology

Electrophysiological measurements were performed 4-6 weeks after AAV infusion. After decapitation, brains were quickly removed and stored in ice-cold partial sucrose solution containing (in mM): sucrose 70, NaCl 70, NaHCO_3_ 25, KCl 2.5, NaH_2_PO_4_ 1.25, CaCl_2_ 1, MgSO_4_ 5, sodium ascorbate 1, sodium pyruvate 3 and D(+)-glucose 25 (carboxygenated with 5% CO_2_/95% O_2_). Coronal slices (300□μm thick) from the hippocampus were prepared in ice-cold partial sucrose solution using a microtome (Leica VT1200S) and immediately transferred in standard aCSF containing (in mM): NaCl 125, NaHCO_3_ 25, KCl 3, NaH_2_PO_4_ 1.2, CaCl_2_ 2, MgSO_4_ 1 and D(+)-glucose 25 (carboxygenated with 5% CO_2_/95% O_2_). Slices placed in the recording chamber were continuously perfused with standard aCSF at 34 °C.

Pyramidal cells on the vicinity of GFP-positive astrocytes were recorded in whole-cell configuration using a Multiclamp 700B amplifier (Molecular Devices, Sunnyvale, CA) and a ITC18 (HEKA) digitizer. Data was recorded using MIES (Allen Brain Institute) written in Igor Pro (Wavemetrics). Somata were patched with borosilicate glass electrodes (Science Products, Germany) with tip resistances of 3-5 MOhm. For sEPSC, cells were recorded at −70□mV holding potential and stable responses were recorded in presence of 10 μM gabazine for 10-15 minutes. The following intracellular solution was used (in mM): Cs Gluconate 130, NaCl 8, HEPES 10, EGTA 0.3, K-phosphocreatine 10, Mg-ATP 4, Na-GTP 0.3, QX-314Cl 1. Synaptic events were detected using Mini Analysis Program (Synaptosoft, Decatur, GA).

AMPAR and NMDAR-mediated EPSCs were evoked by focal electrical synaptic stimulation (0.1 Hz) of the Schaffer collateral pathway, through a borosilicate glass pipette (Science Products, Germany) placed at around 300 μm to the recorded CA1 pyramidal neuron. The intensity of stimulation was set to 10 to 100 μA to produce half-maximal EPSPs. AMPAR-mediated EPSCs were recorded at holding potential of -70 mV for 10-15 minutes in presence of 10 μM gabazine. Intracellular solution for patch pipettes was the same as described above. NMDAR-mediated EPSCs were recorded at + 45 mV for 10-15 minutes with the presence of AMPAR blocker 10 μM DNQX and 10 μM gabazine. A series of consecutive pulses (1, 2, 4, 8) at 100 Hz were given to trigger glutamate spillover(Lozovaya et al., 1999). The interbust interval was 10 s. Synaptic events were analysed using Igor Pro (WaveMetrics). A double exponential model was used to fit the decay kinetics of both AMPAR and NMDAR-mediated EPSCs. To determine the paired pulse ratio (PPR) two pulses were given at 10 Hz. The PPR was calculated by dividing the amplitude of the second peak by the amplitude of the first peak.

### Two-photon glutamate imaging

Mice aged 8-10 weeks were injected bilaterally with 0.7 μl of AAV2/5-GfaABC_1_D::Cas9-HA-Ezr or AAV2/5-GfaABC_1_D::Cas9-HA together with AAV2/5-GfaABC_1_D::SF-iGluSnFR-184V (3:1), as described above (AAV vectors and stereotaxic micro-injections). 4-6 weeks following viral injections, coronal brain slices (350 μm) were obtained as described above (slice preparation and electrophysiology). Bath temperature was set at 32 °C with constant flow (2.5 ml / min) of standard aCSF (in mM): NaCl 125, NaHCO_3_ 25, KCl 3, NaH_2_PO_4_ 1.2, CaCl_2_ 1.3, MgSO_4_ 1, MgCl_2_ 1, sodium pyruvate 3, sodium ascorbate 1 and D(+)-glucose 25, containing CNQX (10μM), APV (50μM) and Gabazine (2μM). Slices were constantly carboxygenated with 5% CO_2_ / 95% O_2_. A custom-built galvanometer-based two-photon laser scanning system was used to image extracellular glutamate (40 x objective; image width, 166.53 μm; excitation wavelength, 900 nm; 100 pixels by 100 pixels per image; acquisition rate, 44.88 Hz). Synaptic release of glutamate was elicited by trains of 10 pulses at high frequency (50 Hz) every 30 s delivered via a borosilicate glass pipette filled with aCSF (60 pA stimulation for 200 μs) placed in the CA1 *stratum radiatum*. Imaging was performed during 6 consecutive trains and subsequently analysed using ImageJ. Fluorescent emission was measured from a circular ROI (diameter, 40 μm) ∽20 μm away from the stimulation pipette. Traces were then averaged and decay tau was calculated by fitting a single exponential function using Igor Pro8 (WaveMetrics). After obtaining baseline recordings, DL-TBOA (10 μM) was added to the bath solution for 10 minutes to block glutamate transporters, and imaging was again performed.

### Contextual fear conditioning

Contextual fear conditioning (CFC) was performed 4-5 weeks after surgery. Mice were handled for two consecutive days prior to conditioning to habituate the animals to the experimenter. CFC was performed in a Plexiglas chamber with a stainless-steel grid floor inside a soundproof cabinet with continuous white noise (68dB; Ugo Basil, Italy). Mice explored the CFC context for 3 min prior to the onset of a foot-shock (0.7 mA, 2 s), and remained in the box for another 30 s before being returned to their home-cage (HC). The CFC context was cleaned with 70% ethanol between each trial. Memory retrieval was performed by a 3 min non-reinforced exposure to the conditioning context, either 5 days (recent) or 28 days (remote) after conditioning. For the novel context group, mice were placed in a context that differed in shape, texture (round, white plastic walls and floor) and smell (2% acetic acid). Automated freezing behaviour was measured using video tracking (Ethovision XT, Noldus, The Netherlands). HC controls received no exposure to the conditioning chamber and remained in their home-cage until they were sacrificed.

### Open field

The open field (OF) was conducted in a white square plastic arena (55 × 60 cm, 25 cm high). Mice were initially placed in one of the corners of the arena and allowed to freely explore it for 10 min. The arena was cleaned with 70% EtOH after each session. Total exploration distance and time spent in the inner circle (35 cm diameter) were measured using the EthoVision tracking software.

### Immunohistochemistry

Animals were deeply anesthetized with Avertine (250 mg / kg) (Sigma-Aldrich Chemie N.V., The Netherlands) and perfused with 4% PFA (Sigma-Aldrich Chemie N.V., The Netherlands) in PBS (pH 7.4). Brains were dissected and postfixed in 4% PFA for 24□h at 4□°C and then transferred to 30% sucrose in 0.1M PBS (pH 7.4). Hippocampal coronal sections (35 μm) were collected serially using a freezing microtome (Leica, Wetzlar, Germany; SM 2000R) and stored in PBS + 0.02% sodium azide. Staining was performed on free-floating brain sections. Sections were blocked with 0.2% (v/v) Triton X-100 (Sigma-Aldrich Chemie N.V., The Netherlands), 2.5% bovine serum albumin (Sigma-Aldrich Chemie N.V., The Netherlands) and 5% normal goat serum (Thermo Fisher Scientific, The Netherlands) in PBS for 1 h. Primary antibodies were diluted in blocking solution and incubated overnight at 4 °C. Sections were rinsed three times with PBS and incubated with secondary antibody diluted in PBS for 2 h at room temperature. Then, sections were washed again three times with PBS and, when needed, incubated with DAPI (300 nmol / l, Thermo Fisher Scientific, The Netherlands) for 15 min at room temperature (RT). Finally, sections were rinsed one more time with PBS and mounted using 0.2% gelatin dissolved in PBS. Sections were coverslipped using Polyvinyl alcohol mounting medium (Sigma-Aldrich Chemie N.V., The Netherlands).

Primary antibodies used: rabbit anti-GFAP (1:1000, Agilent Dako, USA), rabbit anti-Ezrin (1:300, Cell Signalling, The Netherlands), rabbit anti-Iba-1 (1:1000, Abcam, UK), mouse anti-NeuN (1:1000, Millipore, The Netherlands), rat anti-HA (1:500, Roche, Switzerland), rabbit anti-HA (1:1000, Abcam, UK), rat anti-c-Fos (1:500, Synaptic Systems, Germany). All secondary antibodies were used at 1:400 (Thermo Fisher Scientific, The Netherlands): rabbit Alexa 488, rabbit Alexa 568, rabbit Alexa 647, mouse Alexa 647, rat Alexa 488, rat Alexa 568.

### RNA-scope in situ hybridization assay

Mice were injected bilaterally with AAV2/5-GfaABC_1_D::Cas9-HA-Ezr or AAV2/5-GfaABC_1_D::Cas9-HA, in combination with AAV2/5-GfaABC_1_D::eGFP, in order to visualize astrocytes for quantification purposes. Mice were perfused 4-5 weeks after surgery with sterile 4% PFA in PBS and brains were then transferred gradually to 10, 20, 30% sucrose in sterile 0.1M PBS. Brains were embedded (Tissue-Tek, VWR, The Netherlands) and placed in the cryostat at -15 °C for 30 min to equilibrate the tissue before slicing. 10 μm sections from the dorsal hippocampus were mounted on SuperFrost Plus slides (VWR, The Netherlands) and were allowed to dry at -15 °C for 2 h. In situ hybridization with RNA-scope Multiplex Fluorescent Assay was performed following the manufacturer’s instructions (ACD, Biotechne Ltd., UK) (Wang et al., 2012). Briefly, sections were washed with PBS and dehydrated with increasing concentrations of ethanol (50, 70, 100%) for 5 min before being pre-treated with hydrogen peroxide for 10 min. Samples were submerged in target retrieval solution at 99 °C for 5 min, and then treated with Protease III for 30 min at 40 °C. The following probes were used for RNA hybridization (40 °C, 2 h): Ezrin (ACD Bio, Cat. no 535811-C1) and saCas9 (ACD Bio, Cat. no 501621-C3). HRP Signals against each channel (C1 and C3) were then sequentially amplified and developed using TSA Plus fluorophores (Perkin Elmer, The Netherlands) at a dilution of 1:1500, where TSA plus Cy3 (Cat. no. NEL744E001KT) was assigned to C1 probe and TSA plus Fluorescein (Cat. no. NEL741E001KT) to C3 probe. Slices were washed 3 times for 10 min each, followed by IHC that was performed as described above with the following primary and secondary antibodies: chicken anti-GFP (1:500; Abcam, UK) to stain AAV2/5-GfaABC_1_D::eGFP and Alexa 488 anti-chicken (1:400, Thermo Fisher Scientific, The Netherlands). Finally, sections were counterstained with DAPI for 30□s, coverslipped with ProLong Gold Antifade Mountant and dried overnight at RT, in the dark. Ten to 14 images were taken from 5 sections (per mouse) at 40 x magnification. Images were exported to ImageJ to quantify the number of Ezrin mRNA molecules per astrocyte (labelled with GFP).

### Confocal microscopy

A Nikon confocal microscope was used to generate z-stacks (20 μm thick, 1 μm step size) from the CA1 region of the dorsal hippocampus.

For co-labelling experiments (Figure 1), images were imported to ImageJ and merged to form composite images. Then, colocalization of HA with either GFAP, Iba-1 or NeuN was assessed manually using the Cell Counter plugin.

To assess astrocyte morphology (Figure 2), Napa-a (GFP) images were thresholded using ImageJ to remove background signal and, then, astrocyte domain surface was calculated.

For individual counts of c-Fos^+^ cells (Figure 6 and Figure S 6), images were imported into ImageJ and the Cell counter plugin was utilized to mark and count c-Fos^+^ cells within the pyramidal layer of the hippocampus CA1.

### Statistical analysis

Statistical details, number of animals and number of cells are presented in figure legends. All graphs show means ± SEM. Mice with virus misplacements were excluded from analysis. GraphPad Prism 9 software was used for statistical analysis of all data. Outlier removal was performed using the ROUT method (Q=1%). When the data was normally distributed, it was subjected to parametric t-tests and nested t-tests for between groups comparisons. For data not modelled by a normal distribution, the Mann-Whitney U test was used. In case of comparisons that involved more than two groups, analyses were performed by a Two-way ANOVA followed by a post-hoc Bonferroni test. In case of more than two within subject comparisons, a Repeated measures ANOVA was used. All behavioural experiments were replicated twice with independent cohorts of mice.

