## Supplemental Figures for "Proximity of astrocyte leaflets to the synapse determines memory strength"

### Supplementary Figures

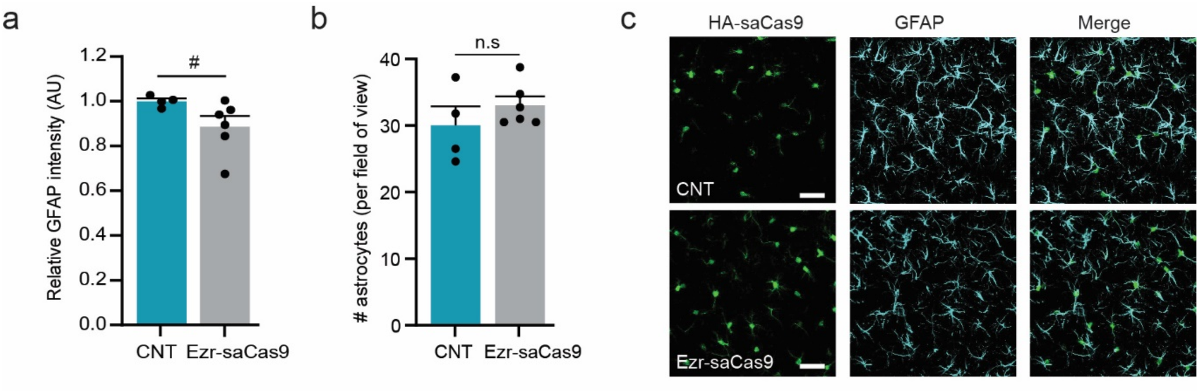


**Supplementary Figure 1. CRISPR-Cas9-mediated depletion of Ezrin does not cause astrogliosis and does not reduce the number of astrocytes**. **a)** Summary data of the mean GFAP intensity per astrocyte (relative to control) in the CA1 region of the hippocampus (Control: 6 slices per mouse from 4 mice, Ezr-saCas9: 6 slices per mouse from 6 mice). Unpaired t test: $t_8 = 1.87$, # $p = 0.09$. **b)** Summary data of the number of astrocytes per field of view (Control: 6 slices per mouse from 4 mice, Ezr-saCas9: 6 slices per mouse from 6 mice). Unpaired t test: $t_8 = 1.075$, $p = 0.31$. n.s = not significant. **c)** Representative examples of GFAP protein expression in HA-saCas9 positive astrocytes. Scale bar: 50 µm. Data are presented as mean ± SEM.

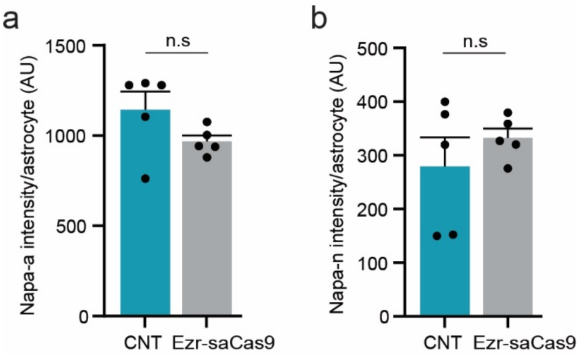


**Supplementary Figure 2. Napa-a (GFP) and Napa-n (mCherry) fluorescence intensities are comparable between groups. A,b)** Quantification of the GFP (A) and mCherry (B) fluorescence intensity per astrocyte (Control: 95 astrocytes from 5 mice, Ezr-saCas9: 144 astrocytes from 5 mice). Unpaired t test: $t_8 = 1.64$, $p = 0.14$ (GFP), $t_8 = 0.92$, $p = 0.38$ (mCherry). n.s = not significant. Data are presented as mean ± SEM.

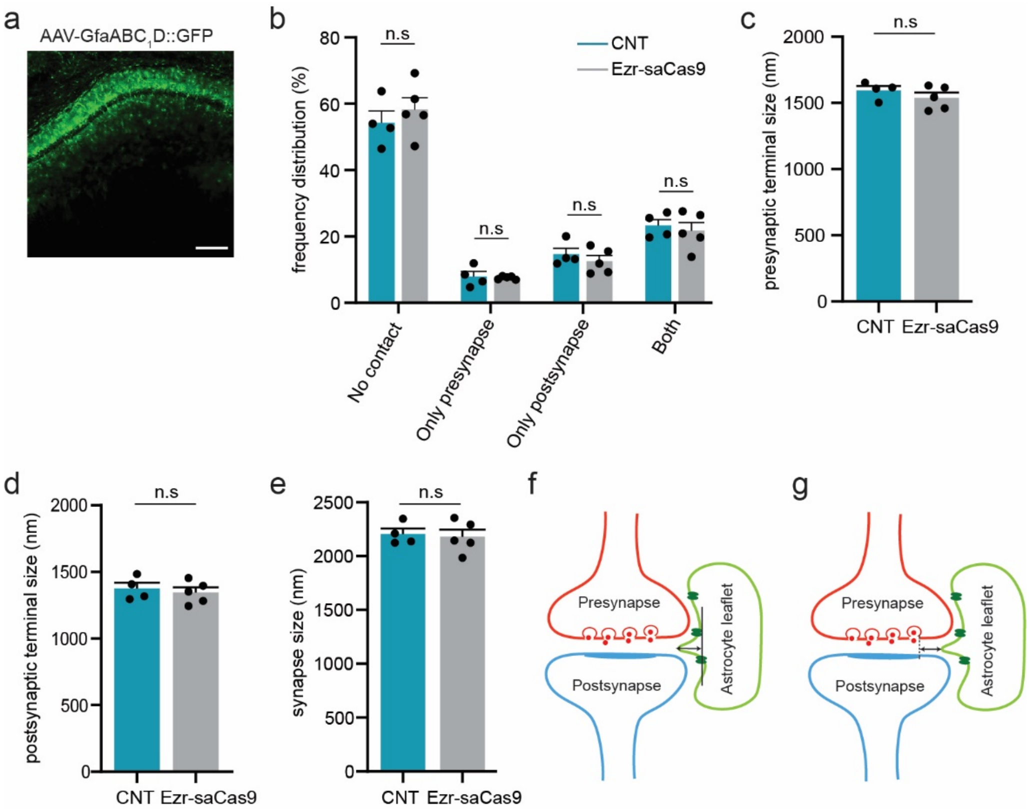


**Supplementary Figure 3. EM analysis of synapse morphology reveals no major changes. a)** Representative image of the AAV- GfaABC$_1$D::GFP expression in the CA1 region of the hippocampus. This virus was infused together with the Control and Ezr-saCas9 in order to confirm sufficient viral expression in samples that were subject to EM analysis. **b)** Summary data of the different types of astrocyte-synapse contact: synapse without an astrocyte contact (no contact), astrocyte contacts only the presynapse (only presynapse), astrocyte contacts only the postsynapse (only postsynapse) and astrocyte contacts both synaptic elements (both). Chi square: $x^2$ (4, 258) = 0.37, p = 0.88. n.s = not significant. **c-e)** Summary data of (**c**) presynapse, (**d**) postsynapse and (**e**) synapse size. Nested t test: $t_{258}$ = 1.48, p = 0.13 (presynaptic terminal), $t_{258}$ = 0.66, p = 0.5 (postsynaptic terminal) and $t_{258}$ = 0.18, p = 0.85 (synapse). n.s = not significant. **f,g)** Example illustrating how the (**f**) astrocyte leaflet length and the (**g**) distance astrocyte leaflet-PSD is measured. Data are presented as mean ± SEM.

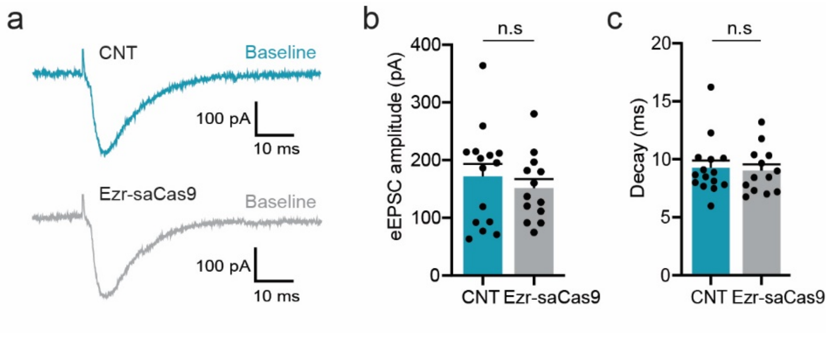


**Supplementary Figure 4. Evoked AMPAR EPSC amplitude and kinetics are not impaired in Ezr-saCas9 mice. a)** Representative traces of evoked AMPAR EPSCs (eEPSCs) from pyramidal neurons from Control and Ezr-saCas9 mice. **b,c)** Summary data of the evoked AMPAR EPSCs (**b**) amplitude and (**c**) decay kinetics (Control: 15 cells from 6 mice, Ezr-saCas9: 13 cells from 6 mice). Unpaired t test: $t_{26}$ = 0.73, p = 0.47 (amplitude), $t_{25}$ = 0.38, p = 0.7 (decay). n.s = not significant. Data are presented as mean ± SEM.

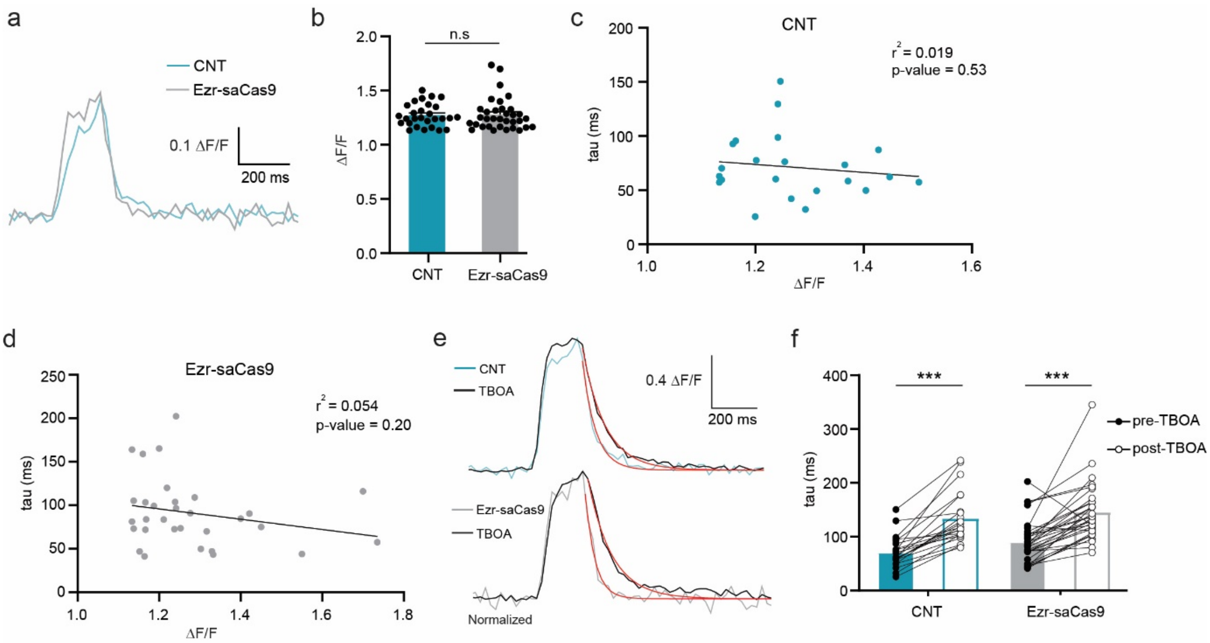


**Supplementary Figure 5. The decay kinetics of the synaptically-evoked iGluSnFR responses do not depend on the magnitude of glutamate release. The iGluSnFR decay kinetics are slower in presence of DL-TBOA in both Control and Ezr-saCas9 mice. a)** Representative traces of synaptically-evoked iGluSnFR response from Control and Ezr-saCas9 mice. **b)** Summary data of synaptically-evoked iGluSnFR responses (Control: 28 astrocytes from 7 mice, Ezr-saCas9: 34

astrocytes from 7 mice). Mann-Whitney test: U = 446, p = 0.67. n.s = not significant. **c,d)** Linear regression plots showing no correlation between iGluSnFR response amplitude and decay kinetics following synaptic stimulation at 50Hz (Control: 22 astrocytes from 7 mice, Ezr-saCas9: 31 astrocytes from 7 mice). **c**) $R^2 = 0.019$, $p = 0.53$ and (**d**) $R^2 = 0.054$, $p = 0.20$. **e)** Representative traces of synaptically-evoked iGluSnFR response before and after DL-TBOA application. Red lines represent the decay kinetics fit for each condition. **f)** Summary data of iGluSnFR decay kinetics before and after DL-TBOA application for Control and Ezr-saCas9 (pre-TBOA and post-TBOA; Control: 28 astrocytes from 7 mice, Ezr-saCas9: 34 from 7 mice). Repeated measures two way ANOVA, treatment effect: $F_{(1,32)} = 86.64$, *** $p < 0.001$. Post-hoc Bonferroni test pre-TBOA vs post-TBOA: (Control) *** $p <0.001$, (Ezr-saCas9) *** $p <0.001$. Data are presented as mean ± SEM.

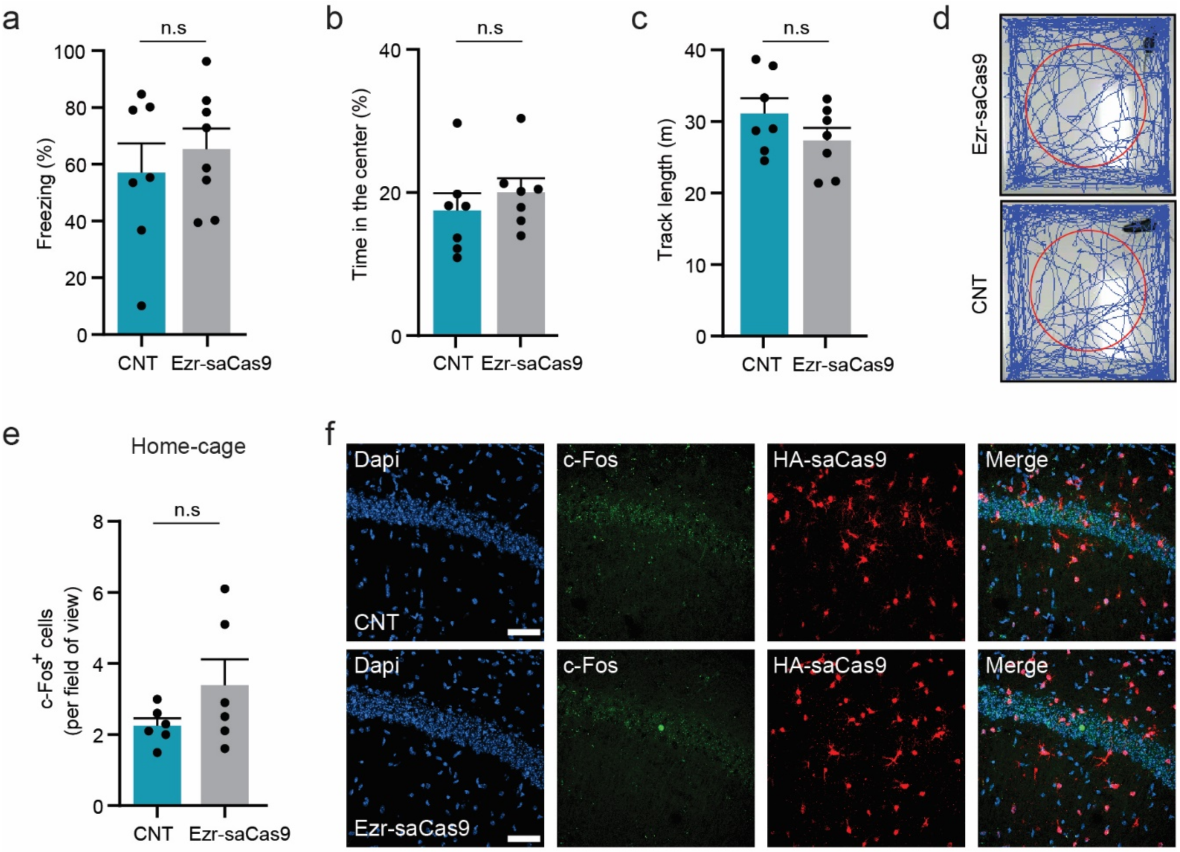


**Supplementary Figure 6. Freezing levels at the remote time point and open field test. Number of c-Fos+ cells from control and Ezr-saCas9 home-caged mice. a)** Summary data of freezing levels assessed at the remote time point show no difference between groups (Control: 7 mice, Ezr-saCas9: 8 mice). Unpaired t test: $t_{13} = 0.67$, $p = 0.51$. n.s = not significant. **b,c)** Summary data of the open field test revealed no effect on anxiety: (**b**) the percentage of time spend in the center of the field and (**c**) the total path length used to explored the field (m) are similar in Control and Ezr-saCas9 mice (Control: 7 mice, Ezr-saCas9: 7 mice). Unpaired t test: $t_{12} = 0.81$, $p = 0.43$ (b); $t_{12} = 1.37$, $p = 0.19$ (c). n.s = not significant. **d)** Representative exploration traces from a Control and Ezr-saCas9 mouse within a 10 min session. **e)** Summary data of the number of c-$Fos^+$ cells within the pyramidal layer of the dorsal hippocampus from home-cage animals (Control: 8-10 slices from 6 mice, Ezr-saCas9: 8-10 slices from 6 mice). Unpaired t test: $t_{10} = 1.48$, $p = 0.16$. n.s = not significant. **f)** Representative images of c-$Fos^+$ cells in the pyramidal layer of the dorsal hippocampus from home-cage mice. Scale bar: 50 µm. Data are presented as mean ± SEM.
